# Aurodox, a polyketide from *Streptomyces goldiniensis*, inhibits transcription of the type III secretion system of multiple Gram-negative pathogens

**DOI:** 10.1101/2024.04.19.589875

**Authors:** David Mark, Nicky O’Boyle, Kabo R Wale, Samantha K Tucker, Rebecca E McHugh, Andrew J Roe

## Abstract

Gram-negative pathogens pose a significant threat due to their propensity for causing various infections, often coupled with formidable resistance to conventional antibiotic treatments. In light of this challenge, the development of antivirulence (AV) compounds emerges as a promising alternative strategy, aiming to disrupt key virulence mechanisms rather than directly targeting bacterial viability. One such compound, aurodox, derived from *Streptomyces goldiniensis*, has exhibited promising AV properties in our prior studies. Specifically, aurodox caused a marked downregulation in the expression and function of the *E. coli* type 3 secretion system (T3SS), a needle-like injectosome structure which is deployed to translocate effector proteins from the cytoplasm to the host target cells.

However, the broader spectrum of aurodox’s efficacy against T3SS across diverse pathogens remained unanswered, prompting the focus of this work. Using quantitative real-time PCR, we show that aurodox exerts inhibitory effects on selected T3SS in various pathogens, including Salmonella typhimurium, *Yersinia pseudotuberculosis*, and *Vibrio parahaemolyticus*. However, aurodox was not a universal blocker of all secretion systems, showing selectivity in its mode-of-action, even within a single strain. This finding was verified using transcriptomics which demonstrated that aurodox selectively blocks the expression of the Salmonella typhimurium SPI-2 type T3SS whilst other pathogenicity islands, including the SPI-1 system were not inhibited. To delve deeper into the mechanisms governing aurodox’s efficacy against these pathogens, we analysed transcriptomic datasets from both *E. coli* and S. Typhimurium treated with aurodox. By identifying orthologous genes exhibiting differential expression in response to aurodox treatment across these pathogens, our study sheds light on the potential mechanisms underlying the action of this rediscovered antibiotic.

## Importance

New treatments to address antibiotic resistance pathogens are urgently needed. Aurodox, a linear polyketide produced by *Streptomyces goldiniensis* has previously been shown to be able to downregulate the expression of a critically important virulence factor, the type three secretion system of *E. coli* and also block host cell colonization. We have explored the wider ability of aurodox to block type three secretion in other species and show that it is capable of blocking the function of T3SSs in further pathogens thereby markedly expanding the known range of pathogens that this compound may be used to combat. This study also shows that aurodox specifically targets SPI-2, a subtype of T3SS crucial for Salmonella persistence within host cells whilst not affecting the SPI-1 system. This finding implies that aurodox likely works through a conserved mechanism and helps reveal insights underlying the action of this rediscovered antibiotic.

Gram-negative pathogens (GNPs) pose a serious threat to public health due to their ability to cause a range of infections whilst exhibiting high levels of resistance to antibiotic therapies^1^. One potential strategy to treat GNPs is the use antivirulence (AV) compounds. The mode of action of these compounds differs from traditional antibiotics as they are not designed to kill or inhibit the infecting organism, but simply to inactivate virulence mechanisms. One such AV target is the type 3 secretion system (T3SS), a needle-like injectosome structure which is deployed to translocate effector proteins from the cytoplasm to the host target cells^2,3^.

Previously, we reported the antivirulence activity of aurodox, a metabolite produced from *Streptomyces goldiniensis*^*4-7*^. In our studies, aurodox abolished T3S in EHEC, EPEC and *Citrobacter rodentium*. As the activity of aurodox in these pathogens is dependent on the inhibition of the master virulence regulator, *ler*^*6*^, we had proposed that only pathogens carrying a LEE-encoded, *ler*-regulated T3SS would display susceptibility to the antivirulence effects of aurodox. To test this hypothesis, we undertook the screening of aurodox against additional GNPs pathogens that carry T3SSs that are phylogenetically distinct and not regulated by *ler*. A range of enteric pathogens encoding T3SSs were selected. These were *Salmonella enterica ssp enterica* serovar Typhimurium, which encodes two distinct T3SS: SPI-1 and SPI-2, *Yersinia pseudotuberculosis* which encodes a Ysc-type T3SS and *Vibrio parahaemolyticus*, which carries two T3SS (VPTTSS1 and VPTTSS2)^2^.

Expression of specific T3SS effectors in each pathogen in response to aurodox was measured using qRT-PCR (Figure 1A-C). In *S*. Typhimurium, we observed downregulation of the SPI-2 effector protein *sseB* (Figure 1A, >23-fold reduction in SPI-2 inducing media, p<0.0001), and marginal upregulation of the expression of the SPI-1 effector *sipC* (0.35-fold change in SPI-1 inducing media, p=0.08). In *Vibrio parahaemolyticus*, aurodox treatment results in a significant reduction in the expression of *vopD* (Figure 1B, >90-fold reduction, p=0.002) however expression of *vopD2* remained unaffected (0.91-fold change, p=0.97). Finally, in *Yersinia pseudotuberculosis*, aurodox downregulated the expression of the Ysc-type effector *yopD* (Figure 1C, >6-fold reduction, p=0.015). As we observed a range of activity across multiple enteric pathogens, these data reveal that aurodox activity is not limited to LEE-encoded, *ler-*regulated T3SSs.

**Figure 1.**
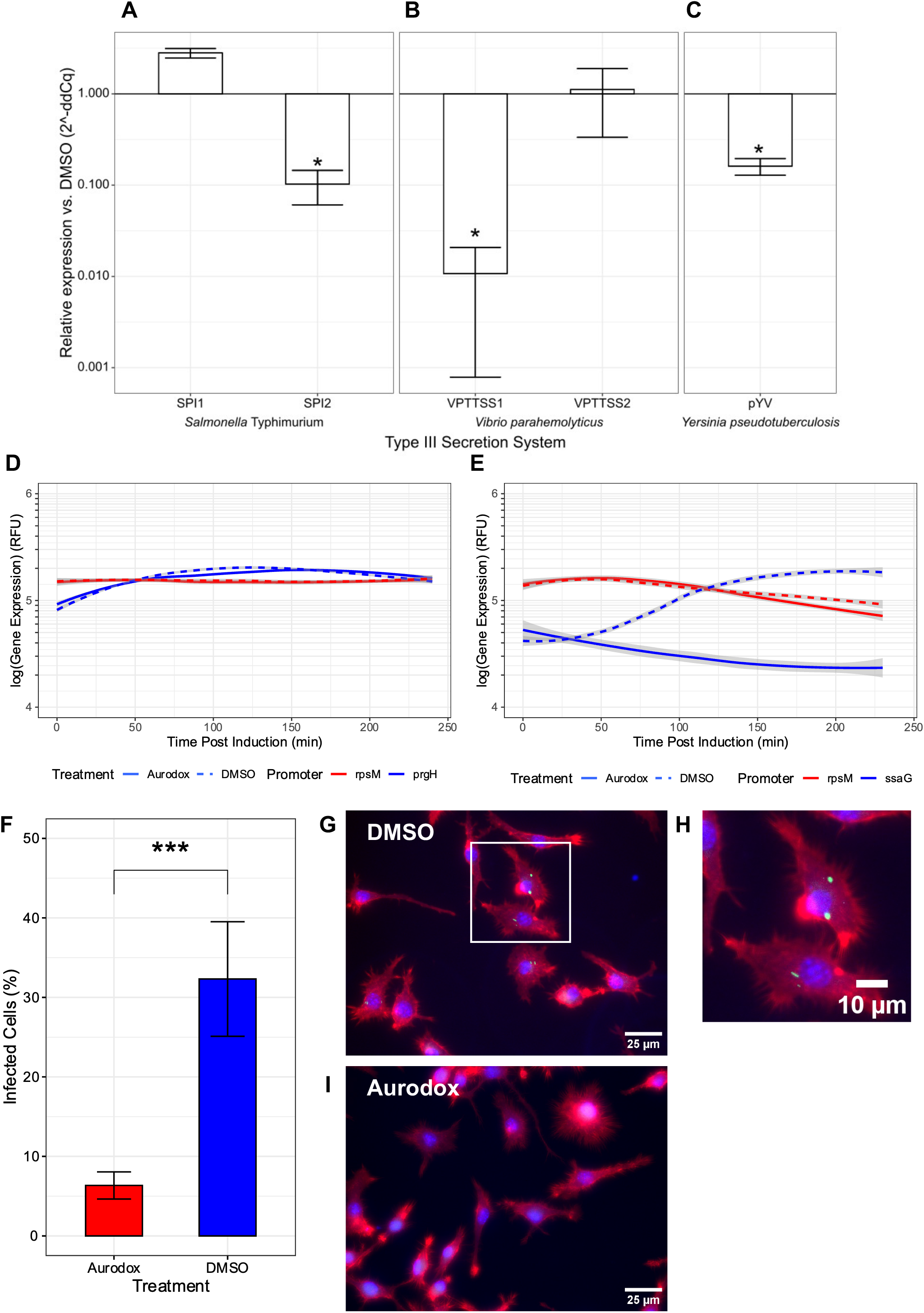
Aurodox inhibitsType III secretion in multiple GNPs. (A-C) Q-RTPCR data show that aurodox inhibited the expression of multiple T3SS effectors including *sseB* of the *Salmonella* Typhimurium SPI-2 TTSS, in SPI-2 inducing conditions (A), *vopD*, secreted by the *Vibrio parahemolyticus* TTSS1 (B), and *yopD* of *Yersinia pseudotuberculosis* (C). It did not, however, repress expression of *vopD2* of the *V. parahemolyticus* TTSS2 or *sipC* of *S*. Typhimurium’s SPI-1; *P<0.05 by One-way ANOVA and Tukey’s post-hoc test. Fig 1D & E: transcriptional reporter assays in *Salmonella* Typhimurium measuring GFP accumulation from the SPI-1 encoded *prgH* promoter (D) or SPI-2 encoded *ssaG* promoter (E). Bacteria were grown to early stationary phase in LB before being switched to inducing media ± aurodox. Aurodox prevented induction of *ssaG* in SPI-2 inducing media, but had no effect on the expression of *prgH*. (F-I) Infection assays showing the protective effect of aurodox against infection of RAW 246.7 Macrophage-like cells by *Salmonella* Typhimurium. After infection in the presence of DMSO, 32% of cells contained *Salmonella*-containing vesicles, compared to only 6% of Aurodox treated cells. (G) Representative image of DMSO treated cell, with SCVs highlighted (H). (I) Representative image of aurodox treated cells.

The differential repression of SPI-1 and SPI-2 in *S*. Typhimurium was further analysed using transcriptional GFP reporter assays^8^. For SPI-1, the expression of the *prgH* promoter (which normally drives expression of a T3SS structural protein) was measured over a six-hour time course (Figure 1D). Similarly, a reporter driven by the *ssaG* promoter was used to measure the transcriptional response of the SPI-2 structural gene to aurodox treatment. Addition of aurodox does not affect *prgH* expression (SPI-1) during the assay, whereas in contrast, there was a marked downregulation of *ssaG* (SPI-2) (Figure 1E). As a control, the ribosomal promoter, *rpsM*, was used for comparison which showed less than 5% variation between aurodox treated and untreated cells. Importantly, the growth of Salmonella was not significantly affected until aurodox concentrations of 8 μg.ml^-1^ were used, which is higher than was required to inhibit SPI-2 (Figure S1). These data demonstrate that the T3SS-inhibitory activity of aurodox is not limited to LEE-encoded, *ler*-regulated T3SSs, and confirms that aurodox has a specific inhibitory effect on the SPI-2 T3SS in *S*. Typhymurium. Moreover, aurodox is not a “universal” blocker of T3SSs.

During Salmonella Typhimurium infection, the SPI-2 T3SS is responsible for maintaining Salmonella within Salmonella-containing vacuoles (SCVs) within intestinal epithelial cells and macrophages. To determine whether the inhibitory effects of aurodox observed for SPI-2 could suppress this activity, the compound was tested in an *in vitro* macrophage model. RAW 267.4 macrophages were infected with S. Typhimurium constitutively expressing GFP to aid visualization. Aurodox treatment significantly reduced the number of RAW cells infected by Salmonella, with 32% of DMSO treated cells infected compared to 6% of aurodox treated cells (>5 fold reduction, p<0.00001) (Figure 1F). Consistent with this change in infection levels, morphological changes associated with Salmonella invasion were reduced in response to aurodox treatment (Figure 1G-I). These results demonstrate that aurodox can exhibit its effect during the intracellular phase of Salmonella pathogenesis, resulting in a reduction in pathogen burden within the macrophage.

The differential effect on SPI-2 over SPI-1 raised the question of how aurodox affects transcription more widely across the genome. To investigate this, whole transcriptome sequencing of aurodox-treated *S*. Typhimurium was carried out. Triplicate cultures were grown in SPI-2-inducing media with either 5 μg/ml aurodox or DMSO. RNA was extracted from each after one hour and converted to cDNA for transcriptomic analysis. Transcripts were mapped to the reference genome and mean fold change and p values calculated. Overall, 11.5% of the genome was differentially expressed in response to aurodox treatment, with 334 genes downregulated and 238 upregulated when compared to the DMSO-treated control (Figure 2A). Differentially expressed genes were identified within the chromosome and three plasmids (pCol1B9, pRSF1010 and pSLT; Figure S2). This analysis revealed that all 32 genes encoded within the SPI-2 pathogenicity island were downregulated in response to aurodox treatment, with the entire pathogenicity island showing statistical significance based on *EDGE test-derived* P value (Figure 2B). These analyses confirm the results of qRT-PCR and GFP-reporter assays as they demonstrate that SPI-2 is downregulated in its entirety in response to aurodox.

**Figure 2.**
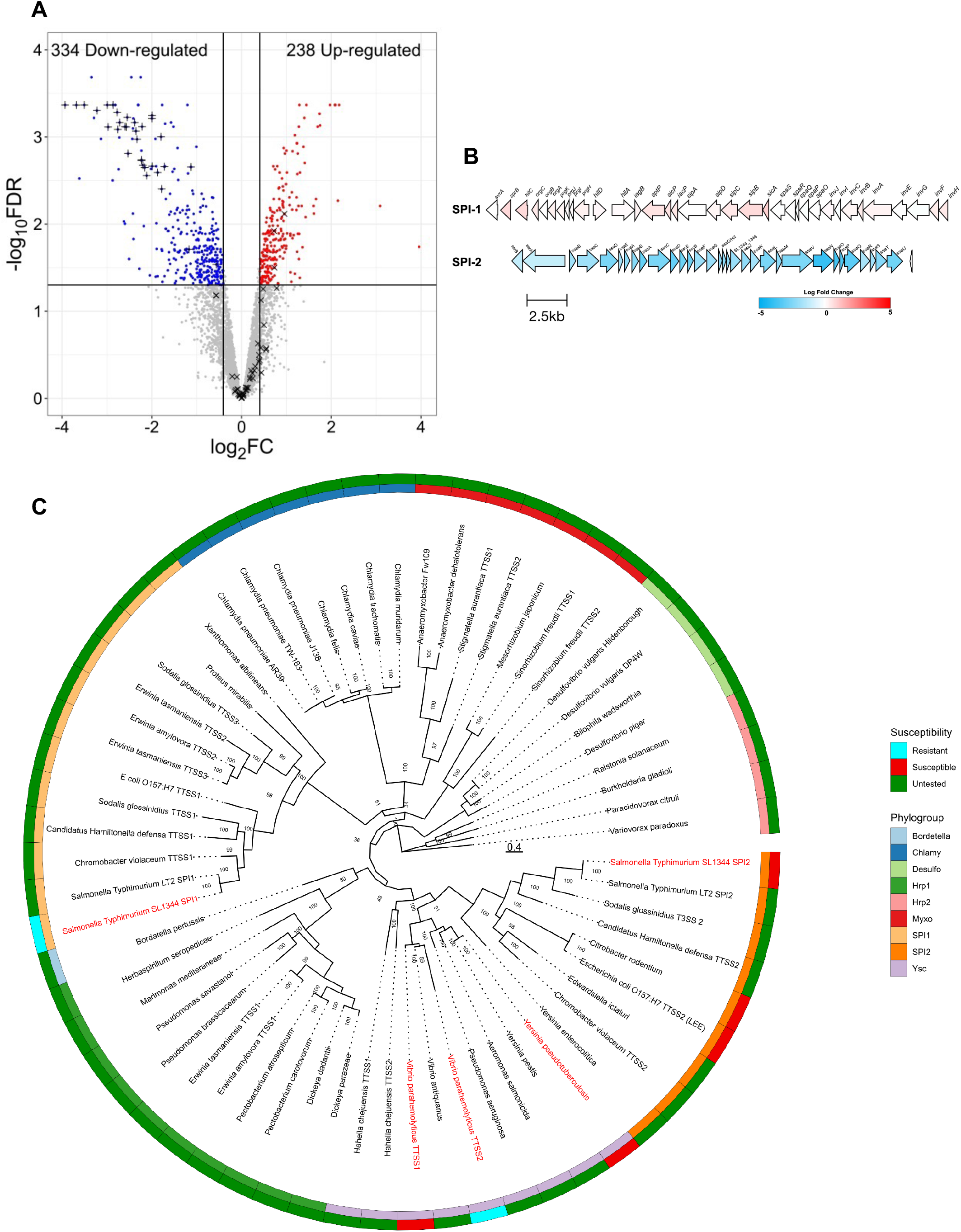
The T3SS inhibiting effects of aurodox may be phylogenetically constrained. (A) Volcano plot showing the global effect of aurodox on the *S*. Typhimurium transcriptome. Genes denoted by a plus (+) are encoded on the SPI-2 pathogenicity island, whilst genes encoded by a cross (×) are encoded on SPI-1 (B) Maps of SPI-1 and SPI-2 illustrating the SPI-2-selectivity in aurodox mediated inhibition of Type III Secretion. (C) Maximum-likelihood phylogenetic tree of selected Type III secretion systems, including those tested against aurodox. We observe that all susceptible T3SSes were placed into a single clade containing the Ysc and SPI-2 phylogroups. Tree generated using IQ-tree, with ModelFinder and 1000 ultrafast bootstraps.

Additionally, gene expression patterns of an additional 12 pathogenicity islands within *S*. Typhimurium were examined in response to aurodox. This revealed that SPI-2 is the only pathogenicity island which is transcriptionally downregulated in response to aurodox treatment (Figure S3). Three genes in SPI 1 *(sicA, sipB, and hilC*) were affected and showed a degree of upregulation.

To gain a clearer understanding of the mode-of-action of aurodox, we identified orthologous genes which were differentially expressed in response to aurodox in both EHEC^6^ and *S*. Typhimurium (Figure S4). From this analysis of the whole transcriptome sequencing 17 orthologous genes were identified, with five genes commonly upregulated and 12 genes downregulated. Several genes which have previously been shown to be involved in virulence regulation were identified, including the alcohol dehydrogenase-encoding gene *adhE*. This finding was of interest because we previously showed that deletion of *adhE* can affect the expression of the T3SS in EHEC^9^. Additionally, multiple amino acid biosynthesis genes were upregulated including the *leu* operon encoding leucine biosynthesis and *met* operon encoding methionine. These analyses identified multiple orthologs which are differentially expressed. To establish a clear link between their altered expression and the observed modulation of virulence will require further experiments.

Of the five T3SSs examined in this study, aurodox was found to inhibit effector expression in three, all of which clustered within the SPI-2 and Ysc phylotypes. In addition, the LEE-encoded T3SSs from EPEC, EHEC and *C. rodentium* which were found to be downregulated by aurodox in our previous study, can also be assigned to the SPI-2 phylogroup^10^. From a phylogenetic tree constructed using core T3SS proteins from multiple GNPs (Figure 2C), we observed that the SPI-2 and Ysc T3SS cluster within one clade. This finding suggests that phylogeny may well be a predictor of aurodox activity. To test this hypothesis, more expansive testing of species within the SPI-2/Ysc clade is required. This should include piscine isolates within this clade, including *Aeromonas salmonicida*^*11*^ and *Edwardiella ictaluri*^*12*^ as they are significant aquacultural pathogens.

## Conclusion

We have shown that aurodox selectively inhibits expression of specific T3SS in different pathogens. In this work we demonstrate activity against Salmonella typhimurium, *Yersinia pseudotuberculosis*, and *Vibrio parahaemolyticus*. However, aurodox does not universally block all T3SS and has a specific inhibitory effect on the SPI-2 T3SS in Salmonella Typhimurium. We also note a phylogroup-dependent activity and a preference for SPI-2 and Ysc phylogroups. The research highlights the potential of aurodox as a promising compound to combat antibiotic-resistant pathogens by targeting virulence mechanisms rather than bacterial viability.

## Acknowledgements

We would like to acknowledge funding from the Wellcome Trust Confidence in Concept scheme for funding work by DM and NOB and the Wellcome Trust Integrated Infection Biology PhD Scheme for funding the PhD of SKT. We like would like to thank the Medical Research Council (MR/V011499/1) for funding REM and AJR. For the purpose of open access, the authors have applied a Creative Commons Attribution (CC BY) license to any Author Accepted Manuscript version arising from this submission.

## Figure legends

**Figure S1:**
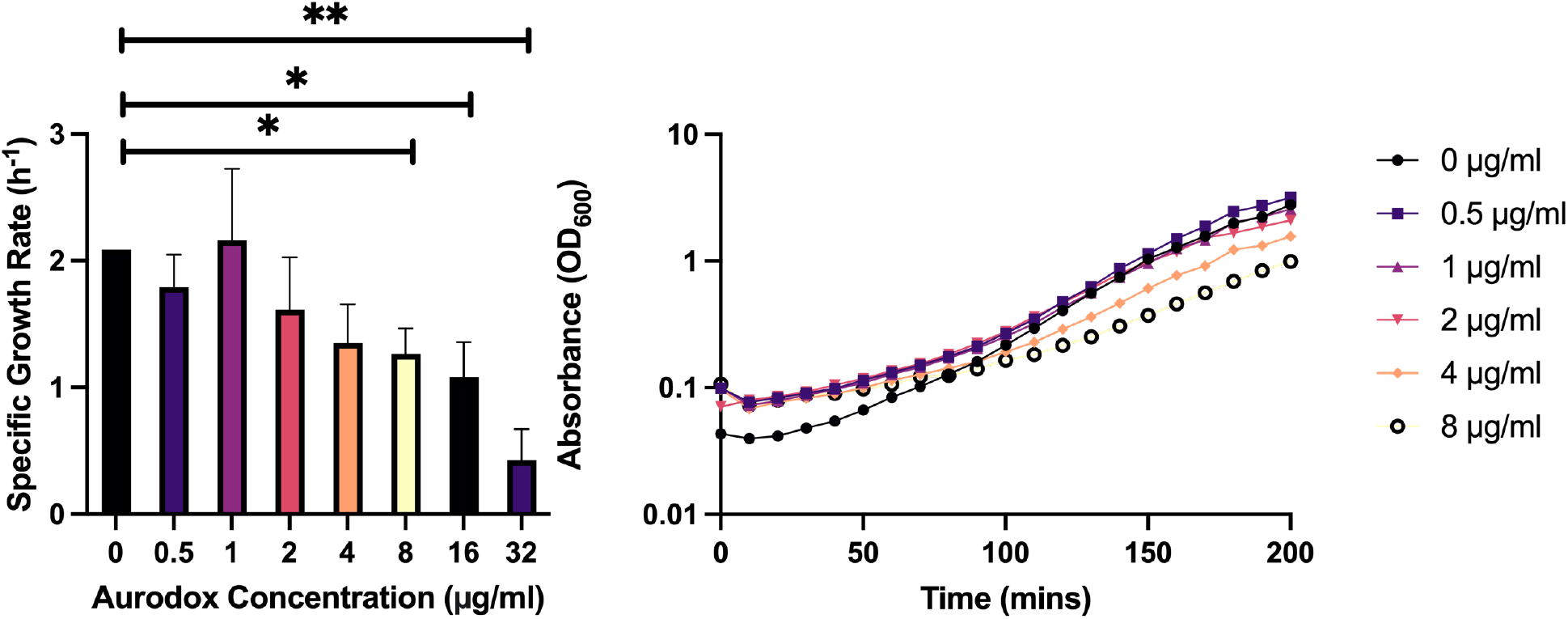
Growth of *Salmonella* Typhimurium is not affected by aurodox.

**Figure S2:**
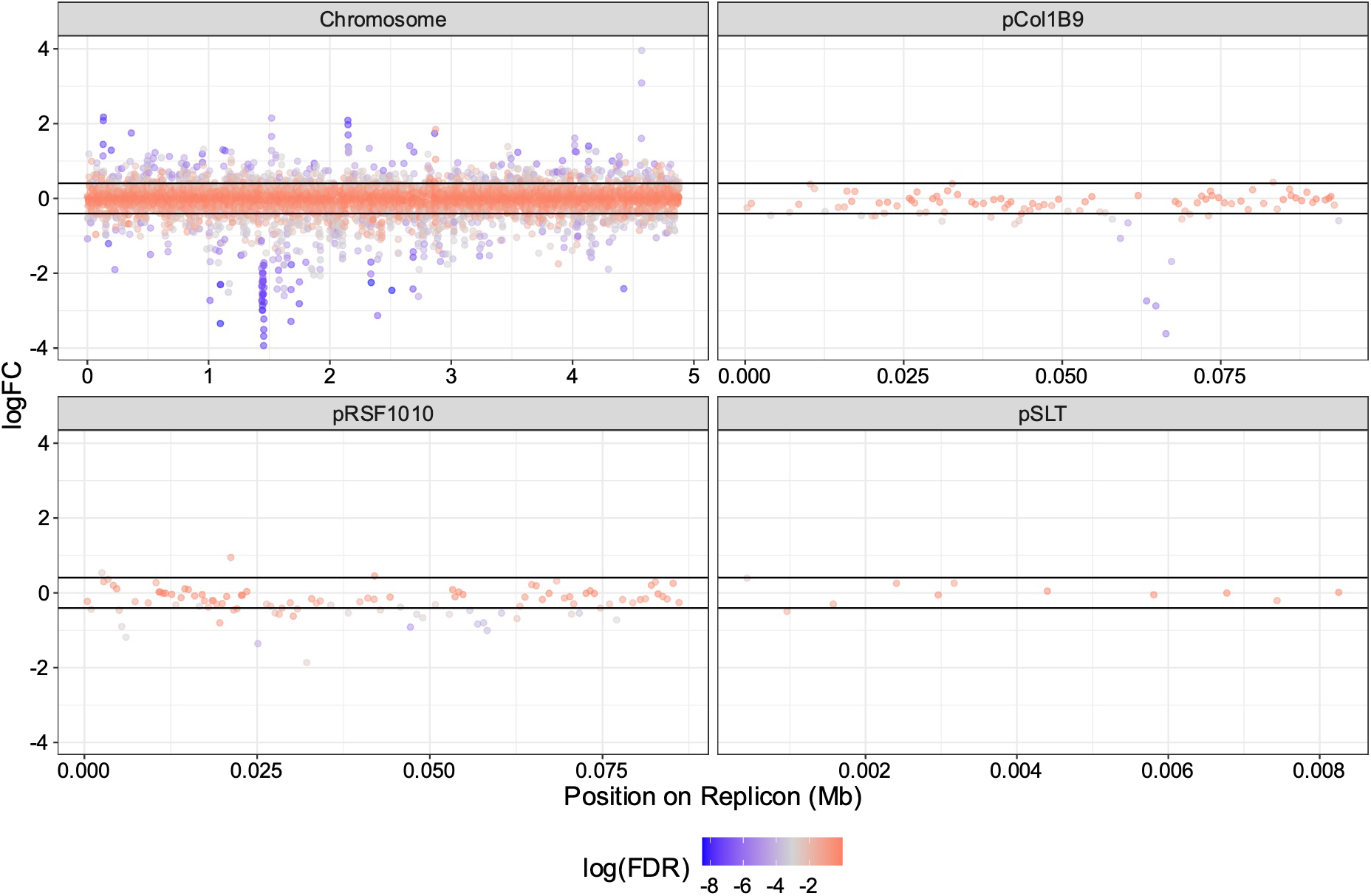
Aurodox affects gene expression at both the chromosome and plasmid level. Map of the *Salmonella* Typhimurium SL1344 replicons. Inhibition of expression was seen on all replicons, except pSLT. We note that expression of SPI-2 effectors encoded on pCol1B9 occurred after aurodox treatment, suggesting polar effects from inhibition of the secretion system.

**Figure S3:**
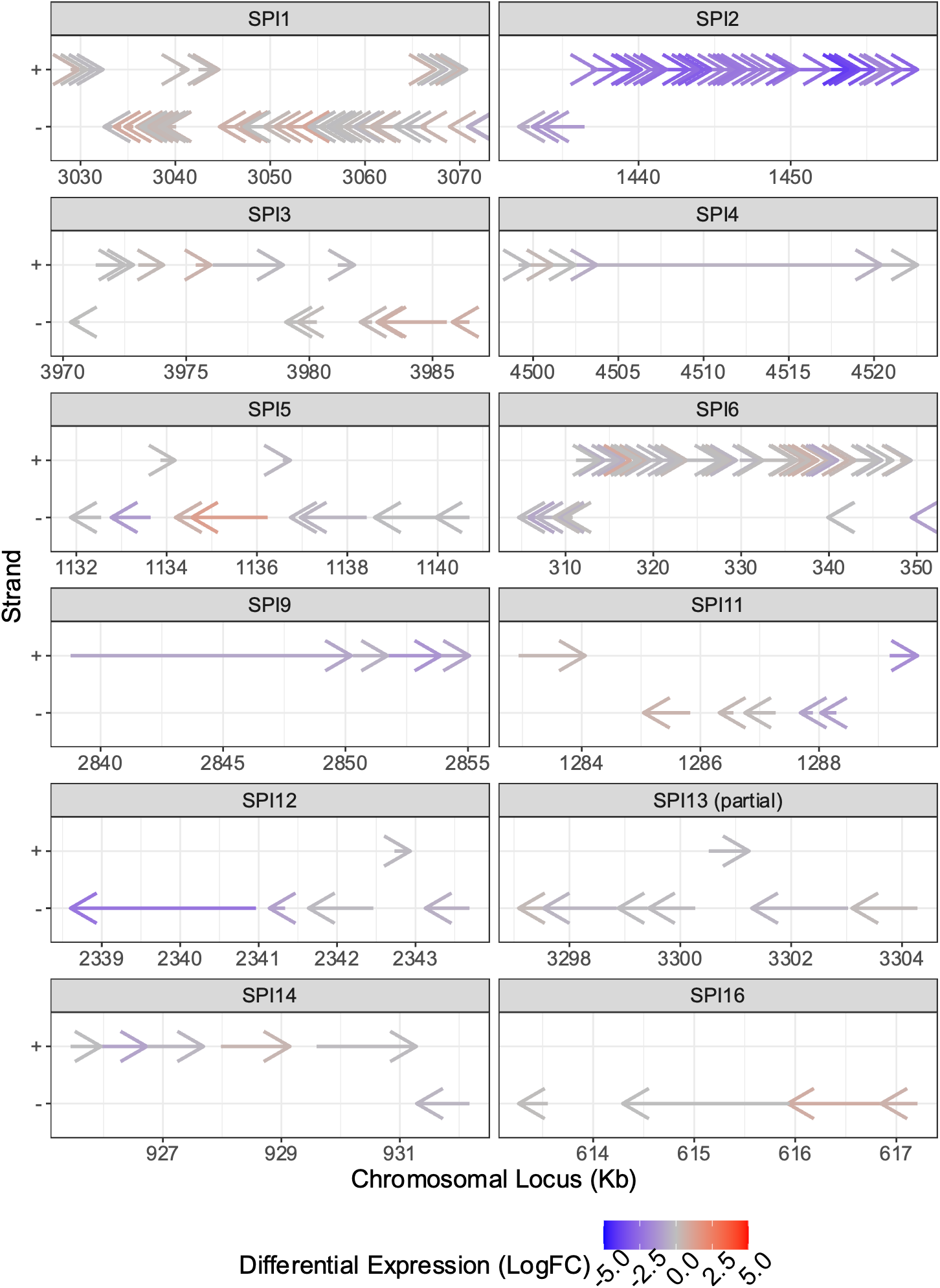
The effect of aurodox on pathogenicity island gene expression is constrained to SPI-2. Maps of the SPI pathogenicity islands encoded on the *Salmonella* Typhimurium chromosome. SPI-2 was the most strongly inhibited island by aurodox treatment.

**Figure S4:**
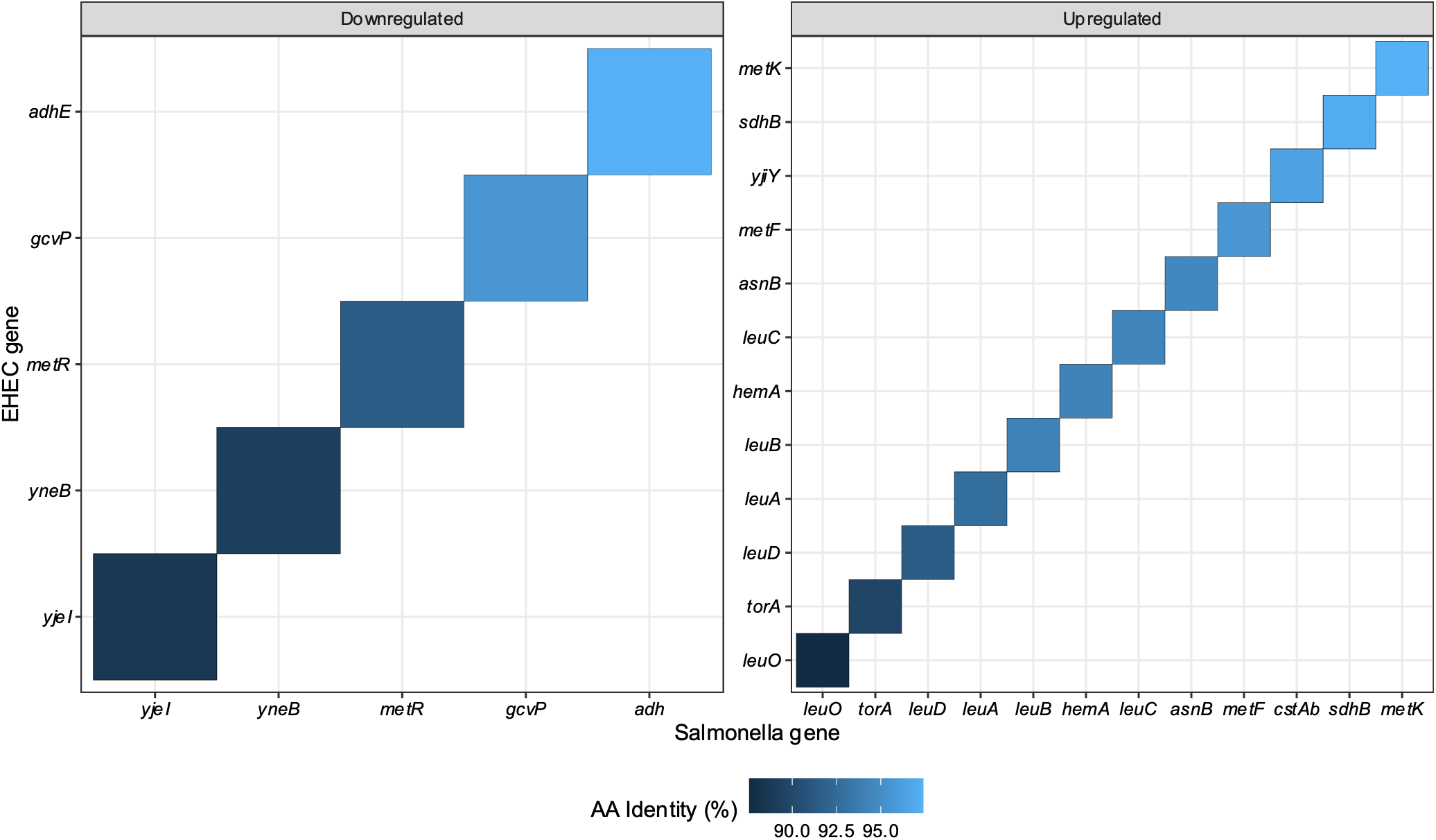
Orthologous genes differentially expressed in EHEC and *Salmonella* Typhimurium post aurodox treatment. Upregulated and downregulated genes identified from EHEC or *S*. Typhimurium in response to aurodox were analysed usingthe BLAST+ Reciprocal best hits tool

## Notes

### Competing Interest Statement

The authors have declared no competing interest.

