## supplementary materials and methods for "Aurodox, a polyketide from *Streptomyces goldiniensis*, inhibits transcription of the type III secretion system of multiple Gram-negative pathogens"

### Acquisition and storage of aurodox.

Aurodox was purchased in its pure form (B.O.C Sciences, Shirley, NJ, USA). A 1 mg $ml^{-1}$ stock solution was prepared by dissolving the compound in dimethyl sulfoxide (DMSO). The overall concentration of DMSO was <0.5% in all experiments.

### Growth of bacterial strains in T3SS-inducing conditions for qRT-PCR

Overnight Lysogeny Broth (LB) cultures were diluted one-hundred-fold in T3SS-inducing media (Supplementary Table 1). Cultures were incubated at 37°C, 200 rpm for 6 h before measuring the OD600 nm and collecting approximately $10^9$ cells by centrifugation at 16,000 × *g* for 2 min.

### mRNA extraction.

Following growth in T3SS inducing conditions (Supplementary Table 3), cells were harvested by centrifugation, resuspended in two volumes of RNA protect reagent (Qiagen, CA, USA) and incubated for 10 mins at room temperature. Following incubation, cells were pelleted by centrifugation at 16,000 × *g* for 10 mins and the supernatant was discarded. Total RNA was extracted by using a PureLink RNA Mini Kit (Thermo Fisher Scientific) according to the manufacturer's instructions after which TurboDNase treatment (Thermo Fisher Scientific) was used to remove contaminating DNA from the samples. For the concentration of extracted RNA, the phenol: chloroform method was used. Briefly, RNA samples were brought to a total volume of 400 µL with nuclease-free water ($NucH_2O$) and then mixed with an equal volume of phenol:chloroform:isoamyline alcohol 25:24:1 (PCIA, Sigma Aldrich). The samples were then centrifuged at room temperature at 13 000 g for 5 minutes, then the upper layer solution was removed and placed in a clean 1.5mL Eppendorf tube. Then 400 µL of chloroform:isolamyl alcohol 24:1 (CIA, Sigma Aldrich) was added to the upper layer in a clean 1.5mL Eppendorf tube, and the tube was vortexed briefly and centrifuged for an additional 1 min at maximum speed. After centrifugation, the upper layer was removed and transferred to a clean Eppendorf tube containing 1 µL GlycoBlue coprecipitate (ThermoFisher Scientific), 40 µL sodium acetate, and

800 µL 100% ethanol. Samples were vortexed gently and stored at −80°C for 1 hour. Following incubation samples were centrifuged at 13 000 g for 20 min at 4 °C. The supernatant was carefully removed to avoid disturbing the pellet. Subsequently, after adding 1 mL of 70% ethanol, the samples were centrifuged for an additional 5 min at 13 000 g. The supernatant was then carefully removed, and the pellet was air-dried. Finally, the pellet was resuspended in 50 µL of Nuc$H_2O$, and the concentration was measured using a NanoDrop DS-11 spectrophotometer (DeNovix).

**qRT-PCR analysis**

Ten nanograms of DNA-free mRNA was used to synthesise 10 µl cDNA with the LunaScript RT Supermix kit (NEB). One microlitre of cDNA was used as a template in qRT-PCR reactions using Luna Universal qPCR Master mix (NEB). The following conditions were employed on a Biorad CFX96 thermocycler: initial denaturation at 95°C for three minutes, followed by 39 cycles of 15 second denaturation and 30 second extension at 60°C. Expression was calculated as $2^{-\Delta\Delta ct}$ relative to expression in DMSO, with *groEL* selected as a housekeeping gene control.

**T3SS phylogeny**

We hypothesized that evolutionary conservation may play a role in our observation that not all tested T3SS were inhibited by aurodox, and so sought to test this by placing them in their phylogenetic context. To do this, we downloaded the protein FASTA data for the strains we tested from NCBI, as well as those of organisms which encode T3SS encompassing previously identified phylogroups (as identified from the secreton database: http://secreton.web.pasteur.fr). These were searched for T3SS using TXSScan as implemented in the Institut Pasteur Galaxy server (https://galaxy.pasteur.fr/). After annotation, the results of TXSScan were manually curated to correct genes that had been assigned to incorrect systems. The amino acid sequences of SctC, SctJ, SctN, SctQ, SctR, SctS, SctT, SctU, and SctV from the remaining T3SS were aligned using ClustalW algorithm as implemented in the msa package in R (v 1.32.0). These alignments were then concatenated, using tools provided by the Biostrings (v. 2.68.1) and tidyverse (v. 2.0.0) packages. Finally, a maximum likelihood tree was constructed using IQ-TREE

(Galaxy Version 2.1.2+galaxy2) with ModelFinder and UF Bootstrap (1000 Bootstraps). The resultant tree was visualised using the treedataverse package and annotated based on T3SS phylogroup and susceptibility to aurodox.

***In vitro* GFP-fusion reporter assays.**

To expand upon our observations that aurodox inhibits SPI-2 but not SPI-1 in *Salmonella* Typhimurium, we employed previously characterised GFP based transcriptional reporters[1,2] (Table S3) To transform *Salmonella* Typhimurium SL1344 with pZEP07, pZEP09, pZEP10, and pAJR70, overnight cultures were diluted 1:100 in fresh LB and grown to an OD (600 nm) of 0.4-0.6. Cultures were then washed thrice in ice-cold 10% glycerol and resuspended in a final volume of 50 µL of 10% glycerol. The cells were mixed with 50-100 ng of plasmid, electroporated at 2.5 kV, and recovered in SOC medium for 1 hr at 37°C before plating on LB + 25 µg $ml^{-1}$ Chloramphenicol (Cm).

For the reporter assays, overnight cultures were diluted 1/100 in 200 µL of LB broth + Cm, and grown in black, clear-bottom plates (Greiner) for four hours (37°C, 200 RPM), with OD (600 nm) and GFP (excitation 485 nm, emission 520 nm) measurements taken every ten minutes. After this, the cultures were centrifuged at 3273 g for ten minutes and resuspended in either SPI-1 or SPI-2 containing media; +Cm, ± 5 µg $mL^{-1}$ aurodox and reincubated as above. Gene expression was calculated as fluorescence intensity divided by OD (600 nm), minus the values from pZEP07.

**RNA-sequencing and associated analysis**

Overnight cultures of *S.* Typhimurium were used to inoculate 10 ml cultures in SPI-2 inducing media (Supplementary Table 3) and grown for 1 hour. Messenger RNA was extracted as previously outlined, RNA quality was assessed by using an Agilent Bioanalyzer 2100 with 100ng $\mu L^{-1}$ concentration accepted as a threshold for individual samples. Library preparation and sequencing was carried out at the University of Glasgow Polyomics facility. Sequencing was carried out on the Illumina NextSeq 500 platform with at least 10 million 100 bp single end reads being obtained.

Quality checking of sequencing reads was performed using FastQC (Galaxy Version 0.73+galaxy0). Following this, reads were aligned to the *Salmonella* Typhimurium genome (Accession GCA_000210855.2) using BWA-MEM2 (Galaxy Version 2.2.1+galaxy0). Featurecounts (Galaxy Version 2.0.1+galaxy2) was used to assign counts to genes, and subsequently differential expression was calculated using edgeR[3] (Galaxy Version 3.36.0+galaxy0), adopting the same cutoff for differential expression as previously[4]. Analysis after this point was carried out in R 4.3.1. To calculate read mapping to SPI2, the getReadCountsFromBAM function in cn.mops[5] was used (window length=20), the read.gff function in ape[6] was used to import annotation data, and further analysis and visualisation was conducted using the tidyverse package.

**Identification of orthologous differentially expressed genes in EHEC and *Salmonella.***

Reads were reanalysed using the same method as described for the *Salmonella* experiment, and subsequently extracted the amino acid sequences of both upregulated and downregulated genes from both datasets. Orthologous proteins in these sets were identified using BLAST RBH (Galaxy Version 0.3.0).

**Infection of RAW 246.7 macrophage with *S.* Typhimurium SL1344**

For the infections, early passage RAW 246.7 cells were seeded onto 13 mm diameter round coverslips, at a density of $5x10^4$ cells $ml^{-1}$ in a 24 well plate (Corning), and incubated for 24 hours at 37$^o$C. *Salmonella* Typhimurium::p*rpsM-gfp* were grown overnight in LB broth + Cm, diluted 1/10 into the same media and incubated at 37$^o$C without shaking until an OD 600 nm of 0.6-0.8. To infect, the *Salmonella* were washed twice in PBS and diluted into RPMI 1640 + Cm ± 5 µg $mL^{-1}$ aurodox or DMSO so as to achieve a concentration of $5x10^6$ CFU $mL^{-1}$, converted using 1 OD (600 nm) = $7.5x10^8$ CFU $mL^{-1}$. Cells were washed twice using PBS, covered in 1 mL of either treated or untreated media, and centrifuged at 400 RPM for 10 mins to associate the bacteria and cells. After one hour, the plate was washed twice with PBS and replaced with clean media containing chloramphenicol, aurodox or DMSO, and gentamicin (30 μg $mL^{-1}$).

After 24 hours, the infections were terminated and stained for imaging – cells were washed twice in sterile PBS, then fixed in 4% paraformaldehyde for 20 minutes at room temperature. After fixation, cells were washed once with PBS and permeabilized with 0.1% Triton X 100 for five minutes, washed, and stained with AlexaFluor555-phalloidin (1:500 dilution, one hour shaking in the dark). After staining, coverslips were mounted in VectaShield mounting medium plus DAPI

Infections were visualised by fluorescent widefield microscopy, at 400x magnification using a Zeiss Axioimager M1 with three infections performed per condition. Images were scored by the proportion of cells in view which were infected and compared using the Wilcoxon ranked-sum test.

**Table S1: Media used in this study**

| **Media Name** | **Components** |
|---|---|
| SPI-1-inducing medium | LB Broth + 5 mM EGTA + 20 mM $MgSO_4$ |
| SPI-2-inducing medium | M9 + 0.4% (w/v) glucose, pH 5.0 (qRT-PCR)<br>M9 + 0.1% (w/v) glucose and 0.004% histidine, pH 5.0 (GFP transcriptional reporter assays and RNA-Seq) |
| *V. parahaemolyticus* TTS1 and TTS2 inducing medium | LB Broth + 5 mM ethylene glycol-bis(β-aminoethyl ether)-N,N,N′,N′-tetraacetic acid (EGTA) + 20 mM $MgSO_4$ |
| *Y. pseudotuberculosis* pYV T3SS inducing medium | LB Broth + 5 mM EGTA + 20 mM $MgSO_4$ |

**Table S2: qRT-PCR primers used in this study**

| Primer Name | Sequence |
|---|---|
| SL1344-groEL-Q-F | GACGCTCGTGTGAAAATGCT |
| SL1344-groEL-Q-R | CCAGAACCACGTTACGGCCT |
| YPIII-groEL-Q-F | GACGCTCGCATTAAAATGCT |
| YPIII-groEL-Q-R | CCAGAACTACGTTACGGCCT |
| RIMD-VopD-Q-F | CGCGGTGAACTGTATGGTTT |
| RIMD-VopD-Q-R | CGTTTGACTCACCGTTGCTT |
| RIMD-VopD2-Q-F | GCGCAACCAGTTATTCCAGA |
| RIMD-VopD2-Q-R | ACATGCTGACTCCACCTTCA |
| YPIII-YopD-Q-F | GCCAGGGAATCAATGTTGCA |
| YPIII-YopD-Q-R | AAACCCATTTCTCGCGCTTT |
| SL1344-SipC-Q-F | GTTCACAGACAGCTTCGCAA |
| SL1344-SipC-Q-R | GTGTAGGACTCAACCCCAGG |
| SL1344-SseB-Q-F | CGAAGGGTATGGTGTTTTGCT |
| SL1344-SseB-Q-R | TCCTGGGTATTTCTGGCACG |
| RIMD-groEL-Q-F | GACGCACGTCAAAAAATGCT |
| RIMD-groEL-Q-R | CAAGCACAACGTTACGGCCT |

**Table S3: Plasmids used in this study**

| Plasmid | Description | Source |
|---|---|---|
| pZep07 | Promoterless *gfp*+, $Cm^R$ | 1 |
| pZep09 | P*ssaG*-*gfp*+ $Cm^R$ | 1 |
| pZep10 | P*prgH*-*gfp*+ $Cm^R$ | 1 |
| pAJR70 | P*rpsM*-*gfp*+ $Cm^R$ | 2 |
